# Structural and dynamic characterization of the C-terminal tail of ErbB2: disordered but not random

**DOI:** 10.1101/2020.09.09.290122

**Authors:** L. Pinet, Y.-H. Wang, C. Deville, E. Lescop, F. Guerlesquin, A. Badache, F. Bontems, N. Morellet, D. Durand, N. Assrir, C. van Heijenoort

## Abstract

ErbB2 (or HER2) is a receptor tyrosine kinase overexpressed in some breast cancers, associated with poor prognosis. Treatments targeting the receptor extracellular and kinase domains have greatly improved disease outcome in the last twenty years. In parallel, the structures of these domains have been described, enabling better mechanistic understanding of the receptor function and targeted inhibition. However, ErbB2 disordered C-terminal cytoplasmic tail (CtErbB2) remains very poorly characterized in terms of structure, dynamics and detailed functional mechanism. Yet, it is where signal transduction is triggered, via phosphorylation of tyrosine residues, and carried out, via interaction with adaptor proteins. Here we report the first description of ErbB2 disordered tail at atomic resolution, using NMR and SAXS. We show that although no part of CtErbB2 has any stable secondary or tertiary structure, it has around 20% propensity for a N-terminal helix that is suspected to interact with the kinase domain, and many PPII stretches distributed all along the sequence, forming potential SH3 and WW domains binding sites. Moreover, we identified a long-range transient contact involving CtErbB2 termini. These characteristics suggest new potential mechanisms of auto-regulation and protein-protein interaction.

**SIGNIFICANCE:** We report here the first description of the receptor tyrosine kinase ErbB2 disordered tail (CtErbB2) at atomic resolution, using NMR and SAXS. We show that although CtErbB2 exhibits no stable structure, it does exhibit partial secondary and tertiary structures likely important for its function. These structural elements are consistent with an active role of the C-terminal tail in the regulation of the receptor’s activity, thanks to the presence of preformed structures for intramolecular interactions, as well as long-range contacts modulating accessibility of those sites and proline interaction sites distinct from the main tyrosine sites. Together, those results reinforce the view that disordered tails of receptors are more than random anchors for partners.

## INTRODUCTION

The ErbB proteins (ErbB1/EGFR/HER1, ErbB2/HER2/neu, ErbB3/HER3 and ErbB4/HER4) are receptor tyrosine kinases that have been extensively studied following the discovery of their involvement in different types of cancer (1, 2). They are constituted of an extracellular domain involved in ligand binding and receptor dimerization, a transmembrane helix, and an intracellular part made of a tyrosine kinase domain and a C-terminal tail. So far, major steps of ErbB signaling have been identified: ligand binding to ErbB1, ErbB3 or ErbB4 induces a conformational rearrangement of their extracellular domain from a tethered into an “open” form that is poised for homo- or heterodimerization. The kinase domains are activated in the dimers (or potential higher-order oligomers (3)), leading to tyrosine phosphorylation of the C-terminal tails (4). This tail is a hub for phosphorylation-regulated protein-protein interactions with adaptor proteins that trigger downstream signaling, including the RAS/MAPK, PI3K/Akt, Src kinases and STAT transcription factors dependent pathways (4).

ErbB2 has no known ligand, but has a constitutively “open” conformation of its extracellular domain, making it the preferential dimerization partner for other ErbB receptors (5). ErbB2 upregulation is found in up to 20% of breast cancers and correlates with poor prognosis (6), and therefore sparked a lot of efforts for the development of targeting strategies, including monoclonal antibodies binding the extracellular domain and small molecules inhibiting its tyrosine kinase activity (7, 8). These strategies have been widely successful, but are limited by inherent or acquired resistance, calling for new combined therapeutic approaches and dual HER2-targeting (9) that require an extended knowledge of associated molecular mechanisms. While elucidating the structures of the extracellular (10–18), transmembrane (19–21), juxtamembrane (21) and kinase (22, 23) domains has given extensive mechanistic insights, ErbB2 C-terminal tail (CtErbB2) is the only region for which we lack such structural description. It was shown that deletion of this region completely abolishes the transforming potential of the activated receptor, while mutation of its five autophosphorylation sites to phenylalanine reduces it by 92% (24). Each phospho-tyrosine interacts with distinct partners, SH2 and PTB domains-containing proteins or MEMO (4, 25, 26), to trigger different signaling pathways *in vivo* (24, 27). However, the structural features underlying these properties are poorly described. Only a few studies on the C-terminal tail of EGFR, the most studied ErbB receptor, were conducted, and it was suggested that its conformation is dependent on its phosphorylation state and regulates kinase activity (28, 29). Similar mechanisms could be at stake in CtErbB2 function.

We previously performed the NMR assignment of CtErbB2 and demonstrated that it is intrinsically disordered (30), in agreement with predictions based on its sequence composition and with measurements of solvent accessibility (31). Intrinsically disordered proteins (IDPs) are a class of proteins with no major, stable tridimensional structure. Instead, their conformation can be described by an ensemble of interconverting flexible structures. IDPs are over-represented in cell-cycle control and signal transduction (32), most likely due to their specific mechanical properties, their increased capture radius (“fly-casting” mechanism (33, 34)) and their specific but usually low- to medium-affinity binding (35). This binding generally involves short linear motifs (SLiMs) and molecular recognition features (MoRFs) that often fold upon binding, and regulate accessibility through long-range contacts that can be modulated by post-translational modifications. Although the development of drugs inhibiting interactions that involve IDPs is challenging, some strategies are beginning to emerge (36). The description of the features of CtErbB2 unphosphorylated apo-state would be the first step towards a better understanding of the mechanisms that lead to its phosphorylation and interaction with adaptor proteins.

Solution state NMR spectroscopy is the dedicated atomic resolution method for such investigation of conformational heterogeneity and dynamics, and is used extensively for determination of conformational ensembles, combined with small angle X-ray scattering (SAXS) (37). Here we present the first atomic-scale description of CtErbB2, using both NMR and SAXS. We show that although this region is highly disordered, it exhibits partially formed *α* helices, and a long-range contact between its N- and C-termini. Several proline-rich regions also transiently adopt PPII helices including potential SH3- and WW-binding short linear motifs. Together, our data suggest the existence of yet unknown conformational regulation processes of ErbB2 function.

## MATERIALS AND METHODS

### Protein expression and purification

CtErbB2 was expressed and purified as described in our previous study (30). The final construct, depicted in Fig. 1 A, comprises residues 988-1255 of the full-length human ErbB2 receptor (numbered here from 1 to 268), plus four additional N-terminal residues (Gly-Ser-His-Met, numbered from -4 to 0).

**Figure 1:**
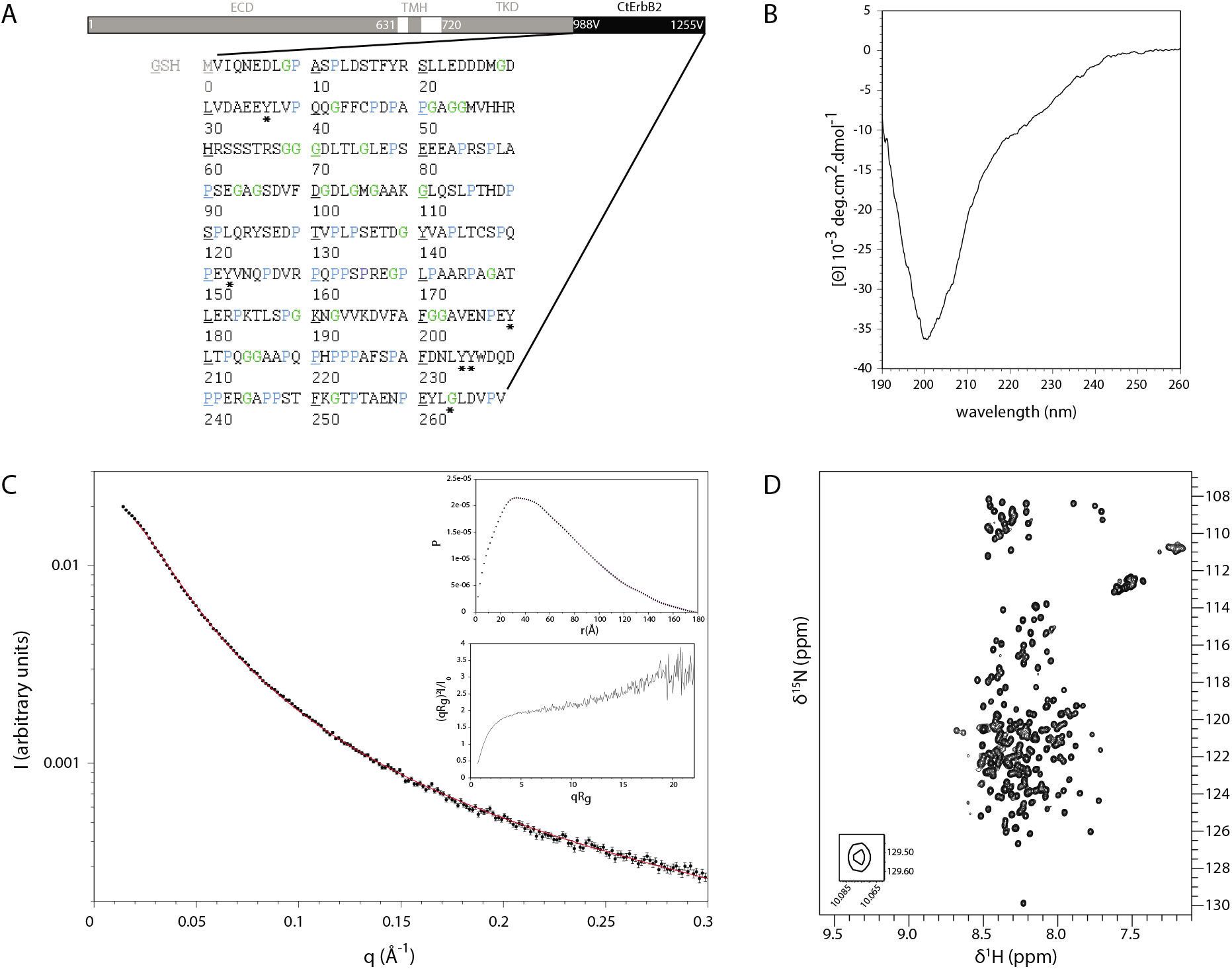
***(A)*** Schematic representation of the human ErbB2 receptor domains and sequence of the CtErbB2 construct. ECD = extracellular domain; TMH = transmembrane helix; TKD = tyrosine kinase domain; CtErbB2 = C-terminal tail. Valine 1 in our CtErbB2 construct corresponds to valine 988 in the full length receptor. Asterisks indicate tyrosines that are phosphorylated by ErbB kinases in dimers of full receptors (autophosphorylation). ***(B)*** CD spectrum (mollar ellipticity per residue) of CtErbB2. ***(C)*** Main Experimental SAXS intensity (black dots) and fit with Sharp Bloomfield equation (red line). *Insets* Top: Distance distribution. Bottom: Kratky plot. ***(D)*** ^1^H-^15^N HSQC spectrum of CtErbB2 at 950 MHz.

### NMR spectroscopy

The typical samples for NMR studies were composed of 200-400 *μ*M protein, in 200-250 *μ*L of MES buffer pH 5.6 (40 mM MES, 200 mM NaCl, 2 mM TCEP, 5% D2O). NMR spectra were recorded at 298 K. Three different spectrometers, each equipped with a TCI cryoprobe (^1^H, ^13^C, ^15^N, ^2^H) and z-axis pulsed field gradients, were used: a Bruker Avance III 950 MHz spectrometer (22.3 T), a Bruker Avance III 800 MHz spectrometer (18.8 T), and a Bruker Avance III 600 MHz spectrometer (14.1 T) (Bruker, Billerica, MA, USA). Data were processed using TOPSPIN 3.5 (Bruker) and analyzed with CCPNMR (38).

#### ^15^N relaxation

^15^N relaxation parameters R_1_, R_2_ and ^15^N-{^1^*H*} steady-state nuclear Overhauser effects (NOEs) of CtErbB2 were measured at 14.1, 18.8 and 22.3 T (600, 800 and 950 MHz proton frequency, respectively), using pseudo-3D HSQC-based pulse sequences comparable to those implemented in the Bruker pulse sequence library. 10 to 14 relaxation delays were used, ranging from 20 ms to 2 s for R_1_ measurements, and from 4 to 450 ms for R_2_ measurements. Relaxation rates were obtained by fitting the decay curves to a two-parameter single exponential decay function. The ^15^N-{^1^*H*} nOes were measured using a TROSY-based pulse sequence using 120° pulses every 5 ms for 4 s for proton saturation. The ^15^N-{^1^*H*} nOe values were calculated as the intensity ratio of peaks in the experiment with or without proton saturation, that were acquired in an interleaved manner.

The ^15^N relaxation parameters were then compared with different models derived from the “model-free” approach (39) and from the Hall and Helfand polymer model (40). Least-square fitting for each residue was done in MATLAB (The MathWorks, Natick, MA, USA) with the trust-region-reflective algorithm. Goodness of fit was assessed calculating reduced *χ*^2^ values for each residue:

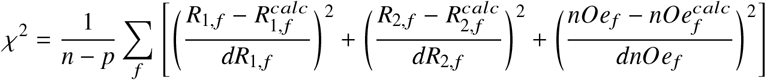

where *dR*_1_, *dR*_2_ and *dnOe* are the errors on the experimental relaxation parameters, n is the number of experimental parameters (here 9, 3 parameters measured at 3 magnetic fields), p the number of fitted parameters in the model, and f the magnetic field. To estimate errors on those fitted parameters, the standard deviation of 1000 Monte-Carlo runs were used. The fit was considered statistically satisfactory if the error function 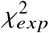 obtained from experimental data (equation above) lay within the 95% confidence limit 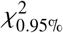 obtained from the 1000 Monte Carlo simulations, i.e. 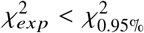.

The model that fitted best the experimental data was the Hall and Helfand model modified by DeJean de la Batie, Lauprêtre and Monnerie (41, 42) (DLM model). The Hall and Helfand polymer model assumes two motional parameters, *τ*_1_ and *τ*_2_, associated with crankshaft-like transitions that consist of segmental motions by rotation around two collinear bonds, and that can be classified according to the relative position in the initial and transition states of the tails attached to the segment undergoing the transition. These transitions can be characterized by the number of conformational jump (s) and the amplitude of the motion of the tails it generates. Transition probability is then determined by the balance between the energy barriers for intermolecular bond rotation and the hydrodynamic forces hindering the motion of segments through the medium. A conformational transition induces distortions of the tails that relax by translations and/or rotations. It was shown from theoretical calculation and Brownian dynamics simulations that these distortions are mostly damped over a limited number of neighboring bonds in polymers (43, 44). The damping process consists of either non-propagative specific motions or distortions of the chain with respect to its most stable conformations. Moreover, it was found that the extent of distortion of the neighboring bonds is often so great as to lead to two simultaneous transitions. This latter process can be considered as a diffusion of bond orientation along the chain. In the case of proteins, correlated transitions correspond to conformational jumps (*ϕ*_*i*−1_, *ψ_i_*) ↔ (*ϕ_i_, ψ*_*i*+1_) ((45, 46). The energy involved in such transitions is controlled by steric interactions (Ramachandran plot), hydrogen bonds and electrostatic interactions. The DLM model additionally takes into account independant libration of the H-N bond, with the introduction of an order parameter A and an associated correlation time *τ_f_*. In our case, the simple Hall and Helfand model did not yield good fit of the data. The addition of the order parameter A allowed a statistically relevant fit, which was not improved by the addition of the third correlation time Tf. The spectral density function was finally:

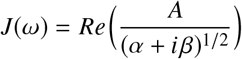

where

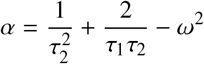

and

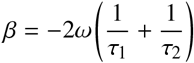

*τ*_1_ is the characteristic time for the correlated bond orientation jumps along the chain while *τ*_2_ describes the damping of the orientation propagation along the chain on each side of the conformational jumps.

#### Residual Dipolar Couplings measurements

The sample was aligned in 6% acrylamide neutral gels. The protein sample was used to rehydrate the gel after dialysis in water and drying at room temperature. The gel was stretched into an open-ended 5mm NMR tube with the help of the apparatus described by Chou *et al.* (47). H_*N*_-N (^1^*D_HN_*) dipolar couplings were measured using a BEST-HSQC-IPAP, at a proton frequency of 800 MHz, at 298 K. The whole procedure was repeated with a different protein sample and a different gel to estimate measurement errors. The respective deuterium quadripolar splittings were 4.98 and 5.66 Hz.

Experimental values were compared with those calculated by Flexible Meccano (48) on two ensembles of 10 000 structures generated without constrains (random coil) or with the following secondary structure propensities: an α helix at 20% between residues 15-20, PPII helices (defined with dihedral angles (−78°, 149°)) at 20% (residues 76-93, 178-191, 216-226 and 238-261), 30% (residues 111-122, 141-151 and 168-175) and 40% (residues 156-165).

#### Scalar couplings measurements

The homonuclear ^3^*J_HNHA_* coupling constant for each residue was measured with the 3D HNHA experiment of Vuister and Bax (49) implemented by Bruker.

#### Paramagnetic Relaxation Enhancement

Four mutants of CtErbB2 were used for four different experiments: C146S, C45S, (C45S,C146S,S227C) and (C45S,C146S,S248C) (for respective paramagnetic tag positions of C45, C146, C227 and C248). These ^15^N-^13^C labeled mutants were reduced with TCEP and incubated at 15°C overnight with the paramagnetic probe (1:10 protein:probe ratio) in a 20 mM sodium borate buffer at pH 8.0. The buffer was then exchanged for the MES buffer pH 5.6 (ZebaSpin Desalting columns, ThermoScientific), removing the excess of free paramagnetic probe. The ^15^N-^1^*H* HSQC spectrum of each paramagnetic form was recorded at 950 MHz. The probe coupling was total, as assessed by MALDI-TOF mass spectrometry (data not shown). To obtain a reference spectrum of the non-paramagnetic proteins, the samples were reduced with ascorbic acid and a second HSQC spectrum was acquired (with peak intensities I_0_ in Fig.2 C). The assignment of each reference spectrum was performed by recording BEST-type HNCA, HNCACB and HNCOCACB (50) and HADAMAC (51) experiments. The probes that were used in this study are 3-(2-Iodoacetamido)-PROXYL (IAP) and S-(1-oxyl-2,2,5,5-tetramethyl-2,5-dihydro-1H-pyrrol-3-yl)methyl methanesulfonothioate (MTSL). The spectra of the probe-attached protein were highly similar to that of the apo-protein for both IAP and MTSL, indicating no major perturbation of the conformational ensemble. Only the results with MTSL, giving similar but stronger effects compared to IAP, are presented here (Fig.2 C).

**Figure 2:**
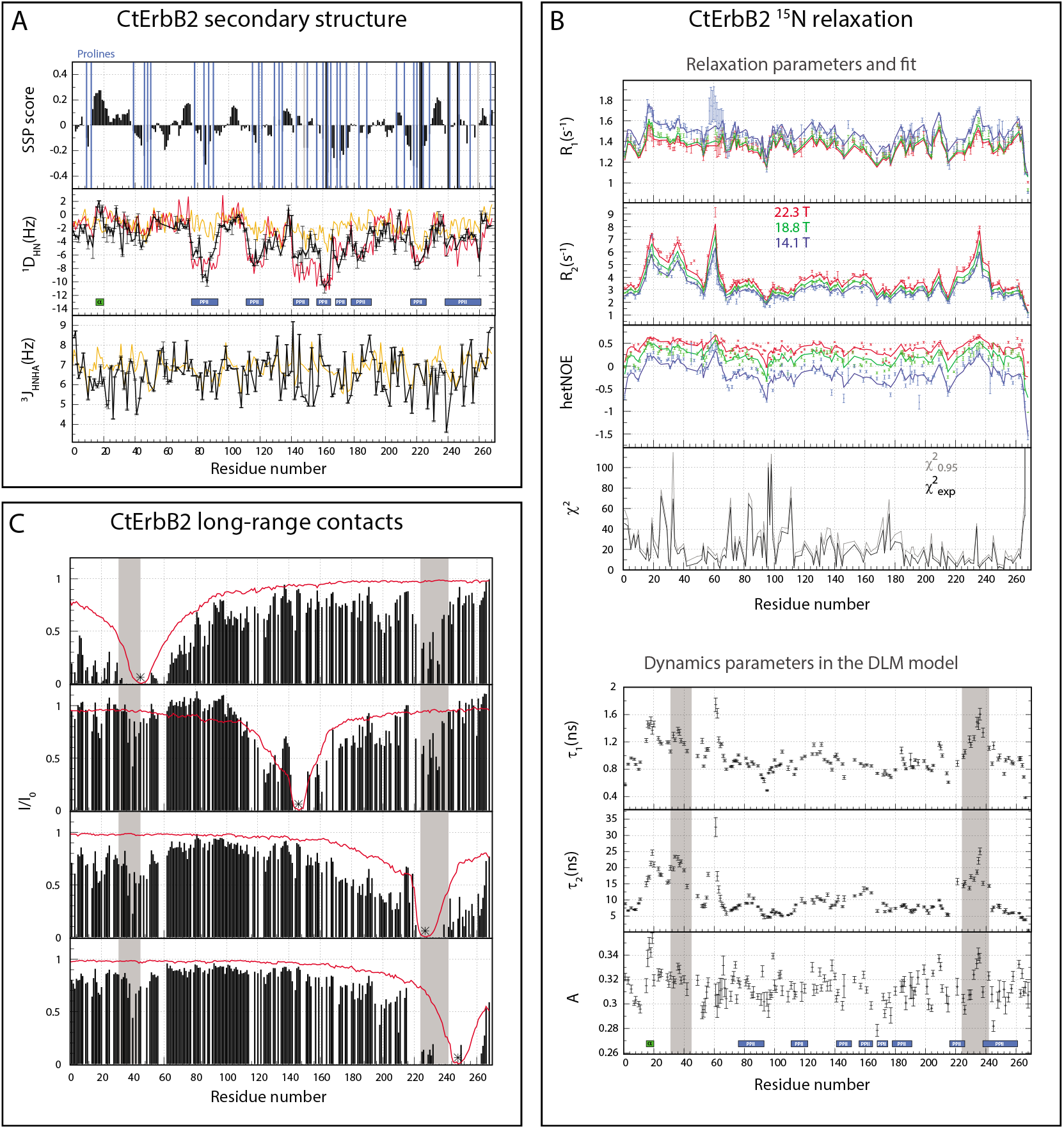
CtErbB2 secondary and tertiary structure identification. Green box= *α* helix; blue box = PPII helix. Grey area: zones of tertiary contact. (***A***) *Top:* SSP scores (58). Vertical bars represent the position of prolines. Blue bars: *trans* prolines (very few or no *cis* conformation detected); Grey bars: prolines with significant proportion of *cis* conformation detected; Black bars: prolines of unknown conformation. *Middle:* N-H residual dipolar couplings compared with Flexible Meccano (48) calculations. Black: experimental; orange: calculated from a random coil Flexible Meccano ensemble (10 000 structures); red: calculated from a random coil Flexible Meccano ensemble with the secondary structure propensities presented below. *Bottom:*^3^ *J_HNHA_*. Black: experimental; orange: calculated for a random coil (63). *(**B**)* Upper pannel: Experimental (points) and recalculated (lines) ^15^***N*** relaxation parameters of CtErbB2 at three magnetic fields using the DLM model described in the Materials and Methods section. *χ*^2^ values were calculated for experimental versus recalculated data 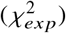 and for 95% lower-*χ*^2^ Monte-Carlo simulations versus recalculated data 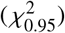; lower pannel: Dynamics parameters of CtErbB2 determined from fitting {^15^*N*}R_1_, {^15^*N*}R_2_ and heteronuclear nOes data at three magnetic fields to the DLM polymer model (see Materials and Methods). Missing data correspond to prolines and weak, unassigned or overlapping peaks. (***C***) Paramagnetic relaxation enhancement measurements for mutants C146S, C45S, (C45S,C146S,S227C) and (C45S,C146S,S248C) of CtErbB2. The paramagnetic probe, indicated by an asterisk, was attached to cysteine 45, 146, 227 and 248, respectively. The red line is the PRE values predicted for the ensemble generated by Flexible Meccano taking into account secondary structures.

### Small angle X-ray scattering

The samples for SAXS were similar in composition to those used for NMR. The measurements were done on a Nanostar Instrument (Bruker) with a Microstar rotating anode (*λ* = 1.54 Å) at 285 K. Several curves were recorded at concentrations from 50 to 200 *μ*M, and were identical, indicating the absence of oligomerization. A Sharp-Bloomfield (52, 53) equation was used to fit the data and obtain polymer parameters for the polypeptide chain (54, 55):

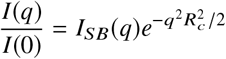

where

- 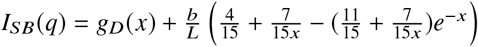
- 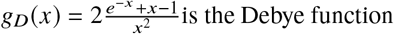
- *x = q*^2^*Lb*/6
- L is the contour length of the chain
- b is the length of the statistical element (twice the persistence length)
- *R_c_* is the radius of gyration of cross section

The theoretical value of the contour length for an protein is calculated as *L = Nl*_0_*f* where N is the number of residues, *l*_0_ is the distance between two sequential C_*α*_, and f is a geometrical factor taking into account dihedral angles. For an unstructured protein we take *l*_0_ = 3.78 Å and f = 0.95 as in (55).

### Circular dichroism

The measurements were performed on a Jasco J-810 spectropolarimeter (Jasco Inc, Hachioji-shi, Tokyo, Japan) equipped with a sample cell temperature control unit (PFD 423S/L Peltier), in a 0.01 mm path length quartz cell. CtErbB2 (200 *μ*M) was examined in the same MES buffer as used for NMR. The wavelength range was 190-260 nm, with a wave step of 0.1 nm. The spectra consisted of an average of ten scans acquired at a speed of 50 nm/min. Spectra were recorded at 298 K. The buffer contribution was subtracted and the signal normalized with protein concentration and number of residues to convert the results to molar ellipticity per residue units (MER).

## RESULTS

### CtErbB2 has no stable local or tertiary structure

The construct of CtErbB2 that we used in this study is presented in Fig. 1 A. Circular dichroism of CtErbB2 was measured at 298 K and shows the characteristic signature of an IDP, with only one strong negative band at 200 nm (Fig.1 B). Small angle X-ray scattering (SAXS) also supports this observation. The scattering curve of CtErbB2 is given in Fig.1 C. The Kratky plot and the distance distribution are characteristic of a disordered protein, giving a radius of gyration of 49.2 Å from the P(r) distribution. Flory theory gives the radius of gyration R_*g*_ of a polymer with N monomers, *R_g_ = R*_0_*N^ν^*, where R_0_ and *ν* depend on the behavior of the polymer in solution, and especially its solvation. Its applicability to denatured or disordered proteins has already been investigated (55). Bernadó and Blackledge (56) found that parameters *R*_0_ = 2.54 Å and *ν* = 0.522 recapitulated well the behavior of such chains. For CtErbB2, this equation gives *R_g_* = 47.4 Å, consistent with our experimental value. It is to be compared with an estimate for a globular protein of the same molecular weight, that would give *R_g_* = 18.7 Å, with 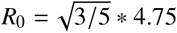 and *ν* = 0.29 (57).

At atomic resolution, the poor dispersion of CtErbB2 proton resonances in the ^1^H-^15^N HSQC spectrum is typical of an intrinsically disordered protein (Fig.1 D). The ^1^H, ^13^C and ^15^N assignment of CtErbB2 (backbone and side chains) was previously achieved (30) and the chemical shifts were deposited into the BMRB (entry 26740). *δ*(^13^C_*α*_), *δ*(^13^C_*β*_) and *δ*(^1^H_*α*_) were used as input for the SSP algorithm (58), and the results are presented in Fig.2 A. The SSP (Secondary Structure Propensity) score is a quantification of secondary structure formation. SSP scores of 0 indicate a fully random coil conformation, while scores of -1 or +1 indicate fully stable secondary structures, either extended (−1) or compact (+1). The SSP scores mainly between -0.2 and 0.2 indicate the lack of fully stable secondary structures in CtErbB2. Moreover, backbone N-H residual dipolar couplings (RDCs) measured in stretched acrylamide gels (^1^*D_NH_*, Fig.2 A) are negative almost all along CtErbB2 sequence, as usually observed for denatured or disordered proteins (59, 60). Overall, all the data show that CtErbB2 is an intrinsically disordered region (IDR).

### CtErbB2 displays local propensities for a residual *α* helix and numerous PPII helices

Many IDPs interact with globular partners through molecular recognition features (MoRFs), which are small sequences exhibiting transient secondary structures in the free state that are stabilized upon interaction. Although these MoRFs can be of all kinds, most of them are *α* helices (61). Given the high content of prolines in CtErbB2, polyproline helices are also likely to be locally populated. We sought to investigate whether CtErbB2 exhibited such transient secondary structures along its sequence, taking advantage of the residue-specific structural information given by NMR parameters, namely SSP scores, H_*N*_-H_*α*_ J couplings (^3^*J_HNHA_*) and ^1^*D_NH_* RDC (Fig.2 A). The deviations of these parameters from expected values for residues in random coil conformation provide information on their propensity to adopt a specific secondary structure.

We firstly investigated the conformation of CtErbB2 numerous prolines. Polyproline helices of type II (PPII) are a common structural motif among disordered peptides and proteins, especially those rich in proline residues (62). These left-handed helices are characterized by an exclusively *trans*-proline conformation, as opposed to the prolines in random coil fragments, for which *cis* and *trans* conformations coexist, with a predominant trans conformation. We previously observed that for all but two prolines in CtErbB2 (indicated by grey bars in Fig.2 A), only *trans* conformation could be identified at our working concentrations (solid blue bars in Fig.2 A), which rules out the possibility of *cis* prolines formed at more than about 10%. To determine whether the absence of peaks for *cis* prolines was due to lack of sensitivity, HSQC spectra at higher concentrations were recorded, which showed many low-intensity peaks, some of which could be assigned to residues following cis prolines (shown in supplementary information, Fig. S1). However, due to still insufficient signal-to-noise ratio, no sequential assignment of these peaks could be achieved.

Typically, positive SSP scores indicate compact structures (3_10_, *α* or PPI helices) which have less than 2 Å of translation per residue, while negative scores indicate extended structures (*β* strands or PPII helices) with a translation per residue typically higher than 3 Å.

In CtErbB2, such negative SSP scores are well correlated with strongly negative ^1^*D_NH_* values, and are predominantly observed in proline-rich stretches (indicated by vertical bars in Fig.2 A). This suggests the presence of PPII helices, rather than *β* strands. H_*N*_-H_*α*_ J-couplings (^3^*J_HNHA_*) make it possible to differentiate these two structures. Indeed, residues involved in *β* strands have higher ^3^*J_HNHA_* (more than 9 Hz) than those involved in PPII helices, which exhibit ^3^*J_HNHA_* around 5-6 Hz, close to or below the random coil value (around 6-7 Hz). In Fig.2 A are presented ^3^*J_HNHA_* values, compared to random coil ^3^*J_HNHA_* predicted for this sequence (https://spin.niddk.nih.gov/bax/nmrserver/rc_3Jhnha/ (63)). Experimental ^3^*J_HNHA_* being almost always equal or below the predicted random coil value, no propensity for *β* strands can be detected in CtErbB2, in extended regions or elsewhere. All our data indicate that the extended, proline-rich regions identified are PPII helices.

Positive SSP scores point to several zones with propensities to form compact helices, that are all N-capped by a serine or aspartate (^14^DSTFYRSLL^22^, ^71^DLTLGL^75^, ^100^DGDLGMG^105^, ^135^SETDGY^140^,^231^DNLYY^235^). The first one is longer and more populated (around 20% according to SSP). It corresponds to higher ^1^*D_NH_* values, consistent with N-H bonds of *α*-helices partially aligned parallel to the gel stretching direction and the magnetic field. These N-capped helices are inserted between extended regions, giving an overall alternation of the two kinds of structures. The same kind of profile was already observed for the measles virus nucleoprotein *N_TAIL_* region (64), in which additionally to the most populated (N-capped) helix, small stretches N-capped by aspartates have positive SSP scores and alternate with more extended stretches.

Globally, all NMR data indicate the presence of transiently populated secondary structures along the sequence of CtErbB2, with a N-terminal *α* helix structured at about 20% and PPII regions with propensities between 20 to 40%. The ^1^*D_NH_* RDCs back-calculated from 10000 conformers generated in the software Flexible Meccano (48) with no secondary structure propensities significantly deviate from the experimental values, whereas thoses obtained using these propensities as input constraints correctly account for the experimental values (orange and red lines in Fig.2 A, respectively).

### ErbB2 exhibits heterogeneous dynamic behavior along its backbone and some level of compaction

NMR, and in particular ^15^N spin relaxation parameters, makes it possible to highlight restriction of dynamics compared to a random coil. The presence of transient secondary structure is expected to locally alter the reorientation dynamics of NH bounds in the 100 ps to nanoseconds timescale that is responsible for ^15^N spin relaxation. Moreover, transient long-range interactions are likely to occur, as part of fly-casting mechanisms (33) and regulation of partner accessibility. The transient modification of the electronic environment due to these long range contacts can also induce chemical exchange and an increase of ^15^N transverse relaxation rates ^15^N R_2_ if they occur in the millisecond time scale.

^15^N R_2_ and R_1_ and {^1^H}-^15^N nOe values, measured at three different magnetic fields (14.1, 18.8 and 22.3T corresponding to 600, 800 and 950 MHz proton frequencies) for CtErbB2, are depicted in the upper pannel of Fig.2 B. The bell-shaped profile that is expected for a completely disordered chain is very distorted, indicating a heterogeneous dynamic behavior, with short fragments exhibiting different type of flexibility (65). To gain more insight into CtErbB2 dynamics behavior, the data were first analyzed using the classically used “model-free” approaches, which didn’t yield statistically relevant fits. Data could be properly fitted with the polymer model developed by DeJean de la Batie, Lauprêtre and Monnerie (41, 42), abbreviated the DLM model. This model, initially designed to describe the dynamics of polymers in solution, appears to be well suited for a long, globally disordered protein chain such as CtErbB2. Our analysis allowed extracting of three dynamic parameters (lower panel of Fig.2 B): an order parameter revealing the amplitude of libration of individual N-H vectors (in a timescale of around 10 ps), *τ*_1_, characteristic of the diffusion of bond orientation along the chain and *τ*_2_, corresponding to the damping of the orientation propagation, i.e. the orientation loss processes. The timescales covered by this approach (tens of ps for reorientation of vectors, low ns local flexibility, and a few to tens of ns when local structure formation is observed) cover the main ranges recently identified as typical for IDPs using the more complex *IMPACT* approach (66).

Along the chain of CtErbB2, low and homogeneous order parameter values A (0.31 ± 0.02) indicate a high proportion of fast, high amplitude, librations motions of NH bonds in the tens of picosecond range, that depends neither on the nature of the residue nor on its implication in a secondary structure. On a slower timescale, the damping of the transitions, indicated by the second correlation time *τ*_2_, exhibit much larger variations. It ranges roughly between 5 and 20 ns (10.4±5 ns), and overall follows the trend of *τ*_1_ (local concerted transitions of bounds orientation), which lies between 0.38 and 1.75 ns (1.00±0.23 ns). Four different types of behaviors can be qualitatively distinguished:

- (i) In the N and C-terminal ends and the glycine rich segment (65-110), *τ*_1_ is around 1 ns or below, typical of a fully random coil, with minimal values of damping correlation times *τ*_2_ around 5 ns. In the middle of the chain, the lowest values of *τ*_2_ are interestingly found near glycines repetitions (*G*_68_*G*_69_*G*_70_, *G*_201_*G*_202_, *G*_214_*G*_215_, or glycine rich segment ([93-110] with 6 glycines).
- (ii) The histidine rich region (56-61) exhibits much more restricted motions with highest *τ*_1_ and *τ*_2_ (almost 2 and 35 ns respectively). Hydrogen bonds can hinder the flexibility of this highly polar stretch, which also contains several arginines and serine.
- (iii) The two segments (15-49) and (225-244) also exhibit high values of both *τ*_1_ and *τ*_2_, indicating a combination of restricted motions at different timescales and fast motions below 1ns. The transient presence of the N-terminal helix can explain this restricted flexibility around the (15-20) segment, with also sightly restricted libration (increased value of A). The other two regions (31-49 and 225-244) display only small helical propensities, and therefore their peculiar dynamic features can not be explained only with secondary structure considerations. Transient tertiary structure (long-range contacts) may be at stake here, that could impeed the damping processes of the bounds orientational transitions. To determine whether this rigidity in the ns timescale was associated with slower dynamics (*μ*s to ms), CPMG experiments were recorded and showed no sign of exchange in this timescale except for rather homogeneous exchange along the sequence attributed to NH exchange with water (data not shown).
- (iv) In the rest of the protein, *τ*_1_ and *τ*_2_ values indicate motions in the low to high ns timescales slightly more restricted than in the glycine rich region. Typically, this encompasses PPII regions, that define dynamically independent stretches separated by more flexible, glycine-containing nodes (green in Fig.1 A). These stretches are often less than ten residues long, not very different from the length of the statistical element estimated for a completely disordered chain (65, 67, 68), explaining the dynamical behavior close to that of a random coil.

SAXS is a method complementary to NMR when it comes to assessing degree of secondary and tertiary structure of a disordered protein (37). To obtain detailed information from the scattering curve, we fitted the intensity to the Sharp-Bloomfield equation, using a polymer model, as previously done for a Repeat-in-ToXin domain (54). This model (red curve in Fig. 1 C) yielded good reproduction of the data with three fitted parameters: the length of the statistical element b ~ 20 Å (which gives a persistence length of 10 Å), the contour length L ~ 770 Å and the radius of gyration of cross section (thickness of the chain) R_*C*_ = 2.7 Å. The 10 Å persistence lenth corresponds to a six amino-acids statistical chain, close to the value of seven estimated by Schwalbe et al.(68) for a chain with few, or short, secondary structure elements. However, the value of the contour length, which reflects the length of the protein when extended without disrupting secondary structures, seems low compared to what is expected for a disordered chain of 272 residues (around 980 Å, see experimental procedures for details of the calculation). This is compatible with the presence of some residual local structure. The presence of many prolines could also contribute to this short contour length, since even low-populated *cis* prolines have an effect, via reduced C_*α*_-C_*α*_ distances.

The effect of structure formation on the radius of gyration is more complex, since the latter can be modified by both secondary and tertiary structures. Comparison of experimental R_*g*_ with those of simulated conformational ensembles is a useful tool to assess the different contributions. Here, the random coil ensemble generated by Flexible Meccano has an average R_*g*_ similar to the experimental one, 50.2 Å. However, when secondary structure propensities are taken into account, and due mainly to the extended nature of PPII helices, this value increases to 57.0 Å, substantially higher than the measured one. This increase in R_*g*_ could be counterbalanced by long-range interactions compacting the protein.

All in all, dynamic studies by NMR and SAXS data show that transient long-range contacts exist in the conformational ensemble of CtErbB2.

### PRE suggests a N-to-C terminal contact

To investigate the existence and location of tertiary contacts in the conformational ensemble of CtErbB2, especially any contact that would explain the specific dynamic behavior of CtErbB2 terminus regions, paramagnetic relaxation enhancement (PRE) was measured. We used four mutants, C146S, C45S, (C45S, C146S, S227C) and (C45S, C146S, S248C), each permetting coupling of MTSL on a single cystein (the two native cysteines and two mutated serines). This mutant collection enables detection of any contact involving the N-terminus, central region or C-terminus of CtErbB2. The *I/I*_0_ values, were *I* is the intensity of each HSQC peak when the probe is paramagnetic, and *I*_0_ when the probe is reduced (diamagnetic) are presented in Fig.2 C, together with the values predicted for the ensemble with secondary structures generated with Flexible Meccano. When the paramagnetic probe is attached to the N-terminus, a decrease in intensity is observed at the C-terminus and *vice-versa*, with a stronger effect when the probe is attached to residue 227 compared to 248. This contact is consistent with high R_2_ observed in the terminal regions, as highlighted in grey in Fig.2 B.

Other contacts are observed between the central part and the extremities of CtErbB2 when the probe is attached at position 146, but they are not observed reciprocally, which could indicate hydrophobic contacts induced by the probe. In agreement with this interpretation is the fact that the region around residues 190-210 is the most hydrophobic in the sequence, and shows paramagnetic relaxation enhancement in all experiments. Globally, these experiments thus strongly suggest a long-range contact between the two segments 30-40 and 220-240 of CtErbB2.

### Conservation of functionally and/or structurally important elements of CtErbB2 amongst mammals

We sought to link our observations on structure and dynamics of CtErbB2 to functional features by looking at the sequence conservation of CtErbB2 amongst mammals. Since conservation of some of the characteristics of CtErbB2 may not require strict sequence conservation but, for example, conservation of high proline density or conservation of interaction motifs, we analyzed sequence conservation node by node on a tree made with mammalian sequences selected with PSI-BLAST. This allowed depiction of more subtle changes, such as displacements of polyproline motifs, compared to analyzing single consensus sequences. The method, as well as the resulting alignments are given in the Supporting Material (supplementary experimental procedures and Fig. S2-S5). The 6 tyrosine autophosphorylation sites are strictly conserved amongst mammals, as expected given their role in signal transduction. Despite the high level of flexibility of CtErbB2, a feature that is often associated with low sequence conservation, the whole region is well conserved amongst mammals even outside these sites (one third of the human sequence is strictly conserved, 15% has a score higher than 0.5 as calculated by Clustal Omega). The same observation was made for EGFR (69). Other structural characteristics of CtErbB2 are then likely to bear functional consequences. Interestingly, the first 50 residues are particularly well conserved (31 are strictly conserved, 10 have scores > 0.5). This region contains the transient N-terminal helix, as well as the N-terminal region involved in long-range transient contacts. The C-terminal region involved in this long-range contact is the second best conserved region (the ^228^PAFDNLYYW^236^ stretch is strictly conserved). Finally, in the PPII regions and PxxP motifs, even though sequence conservation is not strict, proline density is conserved overall, and some mutations of prolines around PxxP sequences compensate each other thus leading to the conservation of the motif (PxxP motifs 6 and 7).

## DISCUSSION

### CtErbB2 disorder is relevant in the context of the whole receptor

All performed experiments indicate that CtErbB2 is mainly devoid of persistent secondary or tertiary structure. CD gives no sign of the presence of stable secondary structure in the protein. Deviation from random coil chemical shifts gives locally no more than 20% of secondary structure, which is rather low even for transiently formed structures in IDPs. Many regions of the protein adopt an extended conformation, but are not drastically constrained in their dynamic behavior. This is consistent with the model of an anisotropic, polymer-like random coil model, where excluded volume effects favor extended conformations (59, 60, 68). This extended nature of disorder gives rise to negative N-H RDCs in stretched gels (59, 60).

Evidence suggest that the disordered behavior observed here is a feature of CtErbB2 in the context of the full-length protein. The other domain that is most likely to interact with CtErB2 is the kinase domain. Keppel *et al.* showed by hydrogen-deuterium exchange followed by mass spectrometry that CtErbB2 is still disordered even in the context of the whole cytoplasmic region (31). Moreover, the disordered nature of CtErbB2 is consistent with its function within the full receptor, which is based on accessibility of interaction sites: its tyrosine side-chains need to be accessible to the kinase domain in the context of activated dimers.

### The N-terminal helix may contribute to regulation of the kinase activity of ErbB2

Despite the disordered nature of CtErbB2, residual helical secondary structure does exist. It is of interest since many IDPs undergo folding upon binding, mostly in *α* helices that are sometimes preformed (70–72). The longest and most populated helix in CtErbB2 is located between residues 14 and 22 (DSTFYRSLL), and contains one tyrosine, which has not been reported to be an interaction site for adaptor proteins *in vivo*. However, in the study by Keppel *et al*. (31), this region was shown to be more solvent-protected than the rest of the C-terminal tail. Furthermore, this helix is visible in the ErbB2 kinase crystal structures, in which it is packed against the kinase domain (PDB 3RCD (23) and 3PP0 (22)). The helix, preformed independently from the kinase, could therefore be stabilized by intramolecular interaction.

Very similar interactions are observed between the kinase and C-terminal tail of EGFR, for which it has been shown that the first dozens of residues of the C-terminal tail have an inhibitory role on the kinase, both from structural and enzymatic data (29, 69, 73) and in terms of transformation potency (74). The involved EGFR helix is called the AP-2 helix, for the FYRAL motif has been shown to interact with the clathrin adaptor protein complex 2 (75). Despite overall moderate sequence conservation between the four ErbB proteins (42.0% similarity maximum, between ErbB2 and EGFR), the region of the helix is well conserved between EGFR (PSPTDSNFYRALMDEED) and ErbB2 (ASPLDSTFYRSLLEDDD), as it is the case for the 50 first residues of the tail (with similarity of 72.5%, see Fig. S5). This could support the hypothesis of a similar kinase-inhibitory role for these residues of CtErbB2. A tyrosine phosphorylation was suggested to disrupt this inhibitory intramolecular interaction in EGFR (29).

### CtErbB2 has long-range contacts involving functionally important regions

The presence of at least one long-range contact is suggested by the relatively small R_*g*_ derived from SAXS compared to an ensemble with the determined secondary structure propensities. PRE experiments also show evidence of a contact between the N- and C- termini, with decreased intensity in one region when the MTSL probe is attached to the other, and vice-versa. Relaxation data confirm that these regions have a particular dynamic behavior. They notably contain less prolines and more hydrophobic and aromatic residues than the rest of the sequence, and are predicted to partially fold in helices (Fig. 2 A). It is not unusual in disordered proteins to observe coincidental secondary and tertiary folding (76). Whether this contact has functional importance remains to be investigated, but its overlap with important sites for signaling is to be noticed. It could affect accessibility of several phosphorylation and interaction sites, such as tyrosine 36 (Y1023 in the full-length protein, the phosphorylation of which was shown to be inhibitory of ErbB2-mediated transformation potency) and tyrosines 234-235 (Y1221-1222 in the full-length protein, an interaction site for Shc) (24). This contact could also be modified by phosphorylation in the activated receptor. In EGFR, an increase in the hydrodynamic radius of the intracellular domain was observed upon phosphorylation (77), correlating with increased dynamics of the C-terminus (29) and disruption of its interaction with the kinase domain (28). Whether CtErbB2 behaves similarly remains to be investigated.

### CtErbB2 contains potential SH3 and WW binding motifs

The PPII (for PolyProline II) conformation is widely spread in proteins, especially in disordered and denatured ones, and in regions containing prolines(62). It was estimated that about 5% of residues (not only prolines) adopt this left-handed, extended triangular helical conformation of 3 residues per turn (78). PPII is still regularly ignored in secondary structure analyses, suggesting an even higher proportion. Given the lack of intra-helix hydrogen bonds and the high solvent accessibility of PPII helices, they are often stabilized by protein-protein interaction. They are mainly involved in interactions with the small SH3 and WW domains. Such interactions can be found in signaling pathways, typically those mediated by receptor tyrosine kinases (79), and WW domains are sometimes considered as the phosphorylation-dependent equivalent of SH3 domains (80). The consensus PPII interaction motif with SH3 domains is PxxP, with +xxPxxP and PxxPx+ (+ being either K or R) defining the most common class I and class II motifs (81). The main WW interaction motifs are PPxY (group I, with an unphosphorylated tyrosine), PPLP (group II), PPR or other proline-rich and K/R containing sequences (group III, potentially in competition for binding with SH3), and phospho-SP or phospho-TP (group IV)(80).

CtErbB2 is strikingly proline-rich (16% of residues are prolines). We could thus expect to find PPII helices in this tail. Negative SSP scores correlated with low ^1^*D_NH_* and^3^ *J_HNHA_* in *trans*-proline rich regions shows that the PPII conformation is indeed sampled at multiple positions along the CtErbB2 sequence. These helices are not especially rigid in the ps-ns timescales, and are only populated to a maximum of 40%, but could adopt a more stable conformation upon interaction. Three PPII stretches contain PxxP motifs: ^81^EEA**P**RS**P**LA**P**SEGA^94^ (2 sequential motifs sharing one proline), ^156^PDVR**P**Q**PP**S**P**REGP^169^ (two nested motifs, RxxPxxP and PxxPxxR), and ^216^AAPQ**P**HP**P**PAFS^227^. The first stretch is highly conserved in mammals (stretch 2-3 in Fig. S2). For the second stretch (number 5-6 in Fig. S2), the 2 nested motifs are not fully conserved, but another PXXP or PXXPXXP motif, with prolines at position 169, 172 and 175, is frequently present in the same PPII rich regions in non human mammalian sequences. Two other PxxP motifs are found:^5^EDLG**P**AS**P**LDST^16^, that is highly conserved among mammals (number 1 in Fig. S2), and ^125^YSED**P**TV**P**LPSE^136^. This last motif is in most non-human sequences part of a PXXPXXP motif with the last proline at position 137 (stretch 4 in Fig. S2). The isolated peptide ^159^RPQPPSPRE^167^ studied by Bornet *et al*. (82) was already shown to adopt a PPII structure, and to be able to interact with FynSH3. One study also suggested that Grb2 SH3 domains bind to ErbB2 (83). Indeed, Grb2 SH2 domain (which is known to anchor Grb2 to CtErbB2 via tyrosine 152) does not recapitulate the binding of full length Grb2 to ErbB2.

There is no consensus PPxY, PPLP or PPR motif in CtErbB2, but the proline-rich regions with R or K residues constitute potential binding sites for group III WW domains. There are also several potential binding sites for group IV WW domains: seven SP motifs (residues 11-12, 86-87, 120-121, 147-148, 164-165, 187-188 and 227-228), and two TP motifs (residues 211-212 ans 253-254). Although a direct interaction between a WW domain and ErbB2 has never been observed, WW-containing proteins such as Pin1 and WWP1 have been shown to have a role in regulation of ErbB2-dependent pathways(84, 85).

CtErbB2 is known to ensure signal transduction thanks to interactions between its phosphorylated tyrosines and SH2/PTB/MEMO domains (4, 25, 26). PPII structures and proline-containing motifs expand the number of possible molecular recognition features of CtErbB2, with potential recognition by SH3 or WW domains.

### CtErbB2 contains two prolines with significant *cis* population

Notable exceptions among prolines are P148 and P259, for which HNCOCACB peaks showing significant *cis* population are detected. However, it is probable that peaks corresponding to a *trans* population are present but superimposed with others, and therefore not assigned. Being located in positions 148 (^146^CS**P**QPEYVN^154^) and 259 (^257^EN**P**EYLG^263^)), the *cis* prolines are close to tyrosines 152 and 261 that were shown to be phosphorylated and important for function (24, 27, 86). Additionally, the phospho-SP sequence of proline 148 could be a binding site for Pin1 prolyl-isomerase, modifying the *cis-trans* ratio. There could therefore be an interplay between proline isomerism and phosphorylation. No sequence or structure determinant can explain the presence of these populated *cis* prolines.

## CONCLUSION

The characteristics of CtErbB2 described here are consistent with several reported functions of the disordered tail in the receptor: disorder ensures high accessibility and adaptability for post-translational modifications and multiple interaction with partners; a long-range contact could modulate accessibility; a preformed *α* helix suggests a regulation mechanism of the kinase activity by the tail; and PPII helices enable possible interactions with SH3 or WW containing proteins. These structural and functional features are summarized in Fig. 3, together with parts of the conformational ensemble generated by Flexible Meccano taking into account secondary structures. CtErbB2 can now be studied in other functionally relevant conditions, investigating in particular how these characteristics are modified by tyrosine phosphorylation, and their impact on the protein interaction network and the regulation of signal transduction.

**Figure 3:**
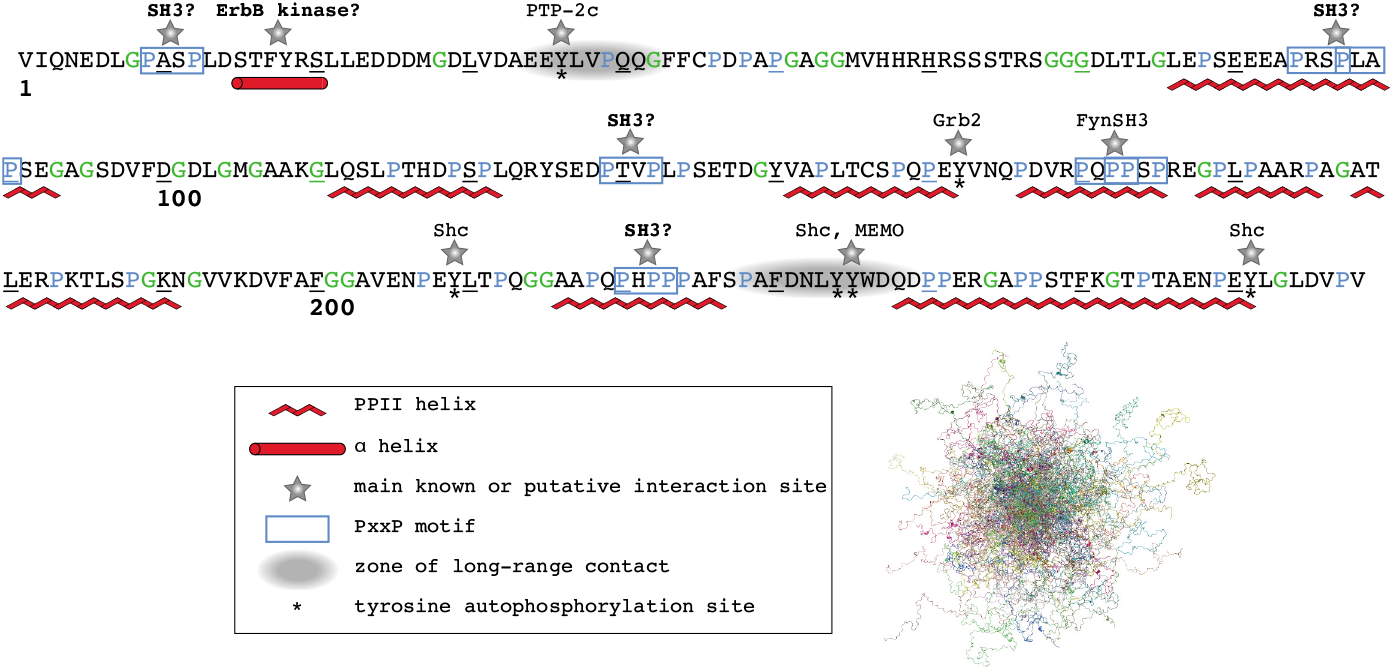
Model of CtErbB2 structural and functional features. *Along the sequence:* The transient secondary structures revealed by SSP scores, residual dipolar couplings and scalar couplings are indicated, as well as the main long-range contact observed in the PRE experiments. In addition to the main known interaction sites and corresponding partners (Shc and Grb2 (24, 25), PTP-2c (25) and MEMO(26) for interaction with phosphotyrosines, FynSH3 (82) for interaction with a RxPxxP motif) the PxxP motifs located in PPII helices are indicated as the best defined putative interaction sites of the proline-rich regions. The putative interaction of the N-terminal helix with the kinase domain is also indicated. *Samples of a conformational ensemble:* Superimposition of 100 structures randomly chosen from the ensemble of 10 000 structures generated by Flexible Meccano with transient secondary structures as input. The structures were aligned on residues -3 to 20.

## AUTHOR CONTRIBUTIONS

Protein expression and purification was optimized and carried out by NA, YHW and LP. NMR experiments were optimized and recorded by LP, YHW, CvH, CD, EL and NM, and analyzed by YHW, LP, CvH and CD. CD experiments were recorded and analyzed by NA and LP. PRE design and experiments were performed by NA, YHW and LP. SAXS experiments were recorded and analyzed by DD. The amino-acid conservation study was performed by FB. CvH, AB and FG conceived the project, CvH, NA and FG supervised research, and CvH managed funding. LP and FG wrote the first draft of the manuscript. All authors discussed results and contributed to the final manuscript.

## ACKNOWLEDGMENTS

We thank Jean-Pierre Le Caer and Vincent Guérineau for mass spectrometry verification of protein purification and probe coupling for PRE measurements, and Aurélien Thureau for help with complementary SAXS measurements. We also thank Arthur Besle and Anaïs Vogel for help in protein preparation and Annie Moretto for buffer, reagents and media preparation. This work was supported by the French National Research Agency (ANR, research grant ANR-13-BSV8-0016 to CvH, FG and AB and financing YW) and the French infrastructure for integrated structural biology (FRISBI). Louise Pinet had a Ph.D grant from the Université Paris-Sud. Financial support from the IR-RMN-THC Fr3050 CNRS for conducting the research is gratefully acknowledged.

## SUPPLEMENTARY MATERIAL

An online supplement to this article can be found by visiting BJ Online at http://www.biophysj.org.

